# Ectopic hepatocyte transplantation cures the pig model of tyrosinemia

**DOI:** 10.1101/648493

**Authors:** Clara T Nicolas, Raymond D Hickey, Kari L Allen, Zeji Du, Caitlin J VanLith, Rebekah M Guthman, Bruce Amiot, Lukkana Suksanpaisan, Bing Han, Maria Giovanna Francipane, Amin Cheikhi, Huailei Jiang, Aditya Bansal, Mukesh K Pandey, Ishan Garg, Val Lowe, Aditya Bhagwate, Daniel O’Brien, Jean-Pierre A Kocher, Timothy R DeGrado, Scott L Nyberg, Robert A Kaiser, Eric Lagasse, Joseph B Lillegard

## Abstract

The effectiveness of cell-based therapies to treat liver failure is limited by the diseased liver environment. Herein we provide preclinical proof-of-concept for the treatment of liver failure through hepatocyte transplantation into lymph nodes in a large-animal model of hereditary tyrosinemia type 1 (HT1), a metabolic liver disease caused by deficiency of fumarylacetoacetate hydrolase (FAH) enzyme. FAH-deficient pigs received autologous hepatocyte transplantation into mesenteric lymph nodes after *ex vivo* transduction with a lentiviral vector carrying the pig *Fah* gene. Hepatocytes showed early (6 hour) and durable (8 month) engraftment in lymph nodes, with reproduction of vascular and hepatic microarchitecture. Subsequently, hepatocytes migrated to and repopulated the native diseased liver. The corrected cells generated enough liver mass to clinically ameliorate disease as early as 97 days post-transplantation, with complete normalization of tyrosine levels and liver function tests. Integration site analysis defined the corrected hepatocytes in the liver as a subpopulation of hepatocytes in the lymph nodes, indicating that the lymph nodes served as a source for healthy hepatocytes to repopulate a diseased liver. Ectopic transplantation of hepatocytes cures the pig model of HT1 and presents a promising approach to the treatment of liver disease in patients with pre-existing liver damage and fibrosis.

**One Sentence Summary:** Transplantation of corrected hepatocytes in mesenteric lymph nodes can cure fatal metabolic liver disease by providing organized liver tissue and by repopulating the diseased liver in the pig tyrosinemia model.

## Introduction

Nearly 14,000 patients wait annually for liver transplantation in the U.S. alone. The problem is considerably worse world-wide and represents one of the most challenging hurdles in medicine.(1) With a universal shortage of organs and limited resources, alternatives to whole organ transplantation are required to address this pandemic. Bioartificial liver devices and repopulation of decellularized liver scaffolds to create bioengineered organs for transplantation have yet to prove effective for the treatment of patients with liver failure. Cell therapy using primary hepatocytes has shown effectiveness in animal models, but the success of this approach has been very limited in the clinical setting.(2) One of the main reasons for this limited success is the inflammation, fibrosis, and scar tissue obstructing blood flow in the failing liver, which constitutes an adverse environment for hepatocyte engraftment and growth.(3)

Hereditary tyrosinemia type 1 (HT1) is an ideal disease model to study treatment options for acute and chronic liver failure in the preclinical environment. HT1 is an inborn error of metabolism of the liver caused by a deficiency of the fumarylacetoacetate hydrolase (FAH) enzyme, which is responsible for the last step of tyrosine catabolism and results in the inability to completely metabolize tyrosine.(4) Untreated, HT1 often leads to fulminant liver failure as early as a few months of life.(5) In the chronic form of the disease, FAH deficiency leads to persistent accumulation of toxic metabolites in the liver, causing oxidative damage and subsequent inflammation, fibrosis, cirrhosis along with high rates of hepatocellular carcinoma (HCC).(6, 7) In HT1, inflammatory changes and liver injury are seen within days to weeks without therapy. Pharmacologic treatment of HT1 exists in the form of 2-(2-nitro-4-trifluoromethylbenzoyl)-1,3-cyclohexanedione (NTBC), a drug which inhibits tyrosine metabolism upstream of FAH, leading to the build-up of less toxic metabolites.(8)

We have previously created and characterized the porcine model of HT1 and showed that this animal is an excellent model of acute and chronic liver failure by reproducing the inflammation, fibrosis and cirrhosis pattern seen in many human liver diseases(9). We have since demonstrated that *ex vivo* gene therapy involving lentiviral transfer of a functional *Fah* cDNA into autologous hepatocytes is curative in both mouse and pig models of HT1.(10) In our previous work, primary hepatocytes were isolated from a partial hepatectomy and transduced *ex vivo* by a lentiviral vector carrying a functional human *Fah* gene. Once corrected, cells were transplanted back into the donor animal via portal vein infusion. However, a substantial clinical limitation to our previous approach is that orthotopic hepatocyte transplantation may not be feasible in patients with acute or chronic liver disease, as the diseased liver is often an inadequate and hostile environment for hepatocyte engraftment and expansion.(11)

Alternative anatomical sites for transplantation of corrected cells could provide a healthier milieu to enable hepatocyte engraftment and proliferation. Lymph nodes are one such alternative site due to several defining characteristics. For example, they are prepared to harbor rapid expansion of T and B cells to support a swift immune response when needed.(12, 13) Lymph nodes are not only capable of accommodating immune cells, but are also a common metastatic site for many types of cancer.(14) They naturally provide a favorable environment for metastatic cell engraftment and growth. This is due, in part, to their high vascularization potential, which permits neoangiogenesis,(15, 16) as well as to their reticular network of fibroblasts and other stromal cells that provide physical and trophic support.(17, 18) Inherent plasticity, together with the fact that their systemic function is not hampered by the transplanted cells,(19) makes lymph nodes a promising site for ectopic cell delivery. Finally, lymph nodes are also easily accessible for both delivery and monitoring of the transplanted cells.

We reported that, after intraperitoneal transplantation of hepatocytes in mice, these cells colonize the lymph nodes and are able to rescue animals from lethal hepatic failure.(20) We explored the mouse lymph node as an ectopic transplantation site for multiple tissues, including liver, and demonstrated that injection of hepatocytes into a single lymph node generated enough ectopic liver mass to rescue the metabolic disorder in the mouse model of HT1.(21) However, it is unknown whether these promising results are translatable into a larger animal model, where a substantially higher hepatic mass would be required for rescue of liver disease. In this preclinical proof of concept study, we demonstrate the therapeutic potential of ectopic transplantation of *ex vivo* corrected hepatocytes into lymph nodes in the HT1 pig, a genetic model of liver failure. In tracking the fate of these cells we show the ability of ectopically transplanted hepatocytes to engraft long term in the mesenteric lymph nodes where they recreate important liver architecture that gives rise to multiple cell lineages seen in the native liver, and serve as a reservoir for repopulation of the recovering native liver.

## Results

### Hepatocytes engraft in mesenteric lymph nodes after ectopic transplantation

To demonstrate that hepatocytes are able to engraft in lymph nodes in a large animal model, a wild-type pig underwent a partial hepatectomy, and harvested hepatocytes were labeled with ^89^Zr (half-life 78.4 h)(22) prior to transplantation into 10-20 mesenteric lymph nodes. Radiolabeling efficiency was ~20% and radioactivity concentration was ~0.1 MBq/10^6^ cells. The animal received 6 x 10^8^ hepatocytes through direct mesenteric lymph node injection. PET-CT imaging at 6 h post-transplantation demonstrated the presence of radioactivity within mesenteric lymph nodes (261.8 ± 108.7 SUV; Fig 1D, Video S1). Radioactivity remained present within the mesenteric lymph nodes at the 54 and 150 h time points (101.1 ± 34.7 and 70.0 ± 26.4 SUV, respectively). Background activity was measured in the left lumbar paraspinal muscle (0.1 SUV). Interestingly, although no radioactivity was detected within the liver at the 6 h time point, increasing amounts of radiotracer were found in the liver at the 54 and 150 h time points (3.3 ± 0.4 and 5.9 ± 0.8 SUV, respectively), indicating accumulation of ^89^Zr-labeled cells or cellular debris. The absence of radioactivity in bone, a known site of uptake for unchelated ^89^Zr, suggested that the radiolabel was retained within the dinitrobenzamide DBN chelator construct and not indicative of unincorporated “free” label. These data at 0.69 and 1.9 half-lives, respectively, suggest possible migration of the transplanted hepatocytes from the lymph nodes to the liver or labeled cellular debris passing through the liver. No radioactivity above background was found in spleen, lung or other organ systems at either of these time points.

**Figure 1.**
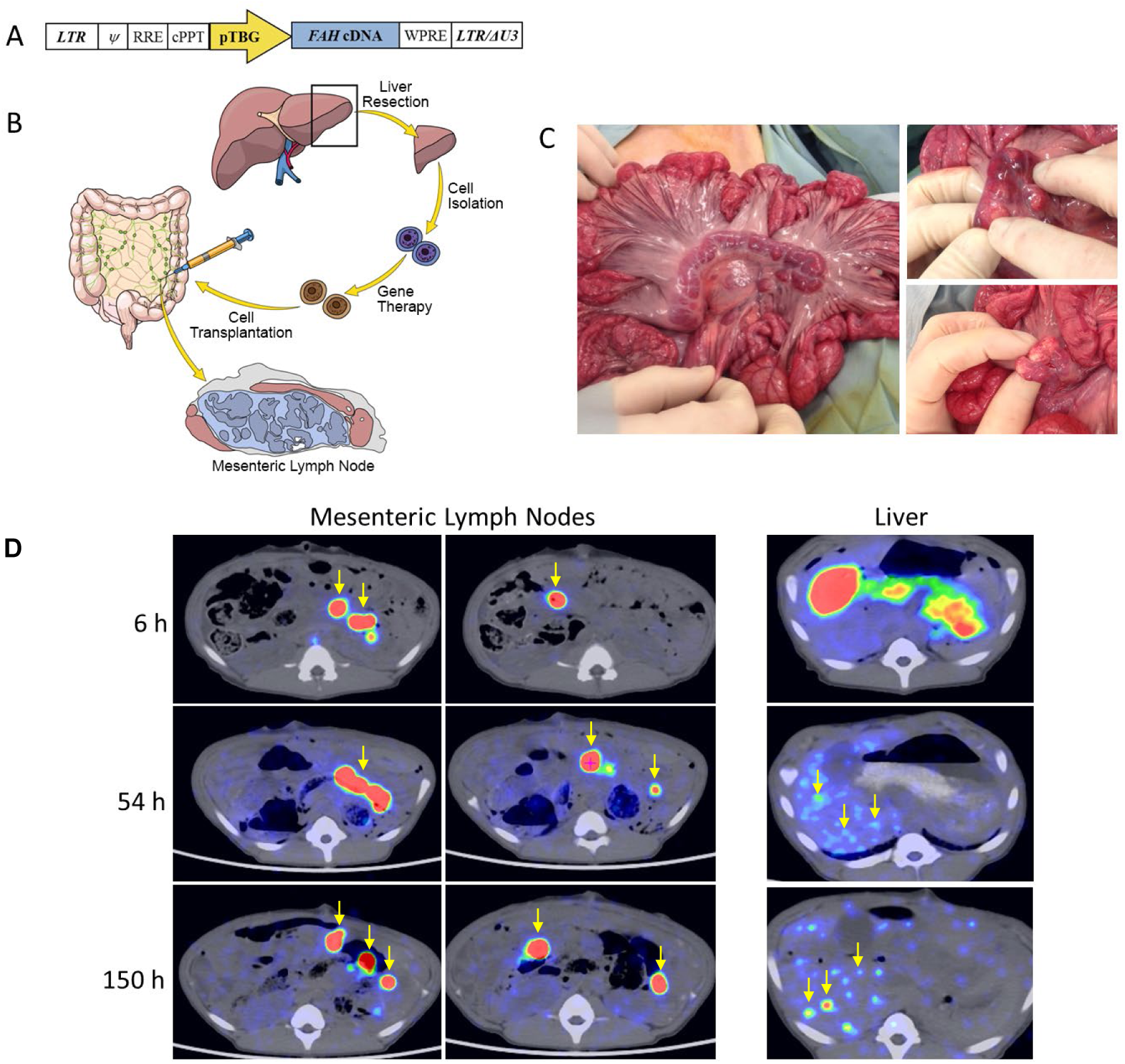
Schematic representation of the lentiviral vector and experimental process, and PET-CT images of 89Zr-labeled hepatocytes at 6, 54, and 150 h post-transplantation into pig mesenteric lymph nodes. (A) The human *FAH* cDNA is under control of the pig thyroxine-binding globulin (pTBG) promoter. LTR, long terminal repeat; Ψ, psi packaging sequence; RRE, Rev-responsive element; cPPT, central polypurine tract; LTR/ΔU3, 3′ long terminal repeat with deletion in U3 region. (B) Four steps are involved in this process: 1) partial hepatectomy, 2) primary hepatocyte isolation through collagenase perfusion, 3) *ex vivo* gene therapy with a lentiviral vector carrying the human *FAH* gene, and 4) transplantation of autologous hepatocytes into the mesenteric lymph nodes. (C) Transplantation of hepatocytes into mesenteric lymph nodes. Macroscopic appearance of the mesenteric lymph nodes before (left) and after (right) hepatocyte transplantation. (D) Representative axial images showing presence of ^89^Zr-labeled hepatocytes in the mesenteric lymph nodes within 6 h of transplantation in a single pig, and engraftment maintained at nearly a week post-transplantation. At the 54 and 150 h time-points, hepatocyte engraftment is observed within the liver as well as in the mesenteric lymph nodes. Representative foci of positive hepatocytes are indicated by yellow arrows (all panels).

#### *Ex vivo* corrected hepatocytes are able to cure a pig model of HT1 after ectopic transplantation into lymph nodes

A total of 5 *Fah*^-/-^ animals were maintained on NTBC until the time of *ex vivo*-corrected autologous hepatocyte transplantation into lymph nodes (Fig 1A-C), at which point administration of NTBC was discontinued to propagate liver injury and stimulate hepatocyte regeneration. This induced selective expansion of the newly transplanted FAH-positive hepatocytes. Animals were cycled on and off NTBC based on weight parameters until NTBC-independent growth was achieved (Fig 2A, Fig S1). NTBC-independent growth was attained at a mean of 135 ± 25 days post-transplantation (range: 97 to 161 days), after 3 to 6 cycles of NTBC.

**Figure 2.**
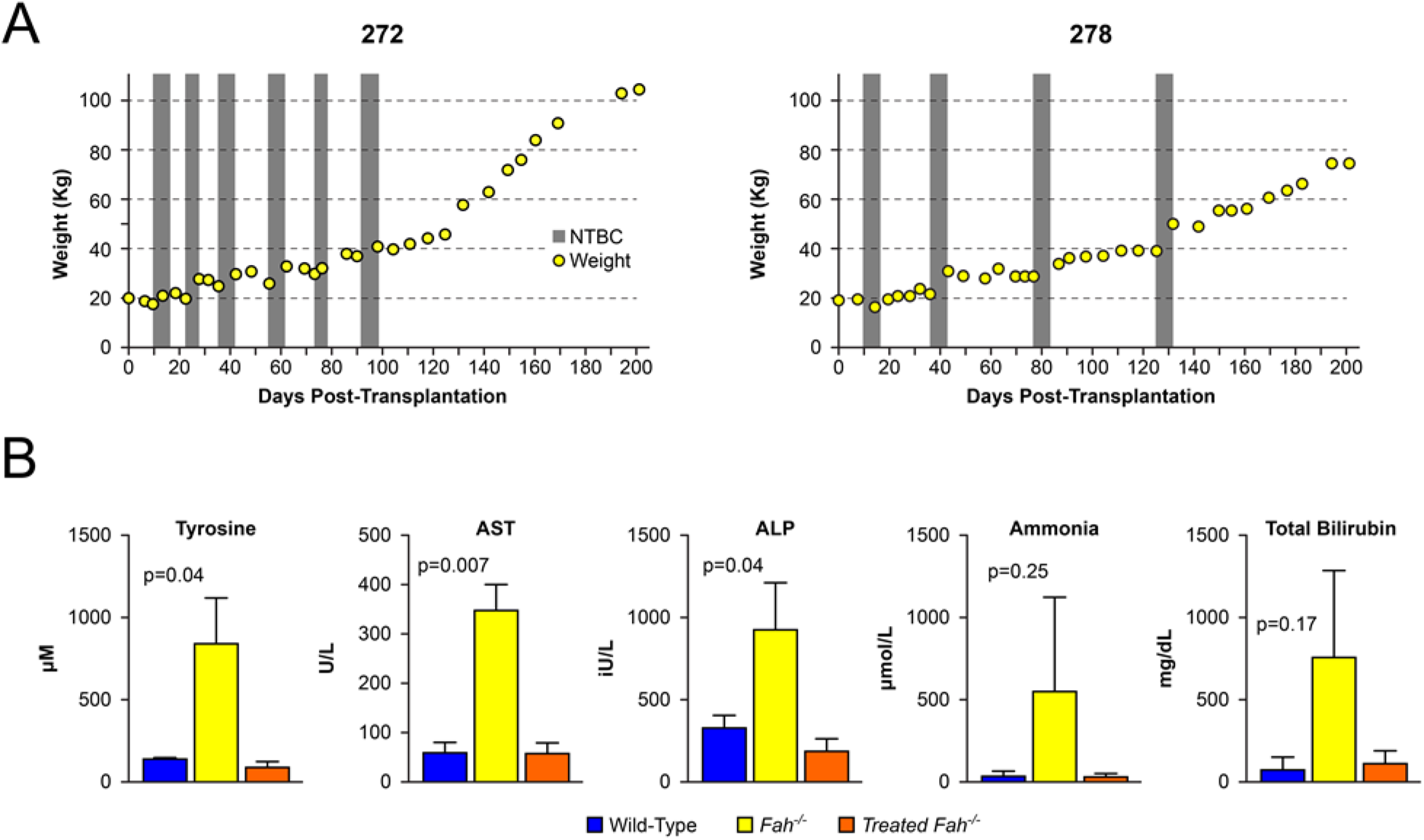
Weight gain and biochemical correction of treated pigs. (A) Weight stabilization of pigs 272 and 278 demonstrating NTBC-independent growth at days 97 and 131 post-transplantation. (B) Normalization of tyrosine, aspartate aminotransferase (AST), alkaline phosphatase (ALP), ammonia, and total bilirubin levels at the time of euthanization in all animals. Mean ± SD are shown for experimental animal cohort (treated *Fah*^-/-^ animals) alongside historical wild-type and untreated *Fah*^-/-^ control animals. P-values are provided for experimental animal cohort compared to untreated *Fah*^-/-^ controls.

Biochemical cure of the HT1 phenotype was confirmed by normalization of liver specific enzymes known to be elevated in HT1 as well as normalization of tyrosine levels. At the time of euthanasia, mean tyrosine levels (84.2 ± 34.5 µM) in the five treated pigs were within normal limits for wild-type animals, and were significantly lower than untreated *Fah*^-/-^ controls (826.3 ± 277.5 µM) (Fig 2B). Similar results were seen in liver function tests in treated animals (AST: 56.8 ± 21.7 U/L; ALP: 186.6 ± 78.8 IU/L; ammonia: 35.8 ± 14.4 µmol/L; albumin: 3.6 ± 0.3g/dL; and total bilirubin: 0.15 ± 0.1 mg/dL) compared to untreated *Fah^-/-^* controls (AST: 343.7 ±54.5 U/L; ALP: 918.3 ± 282.5 IU/L; ammonia: 553 ± 561.3 µmol/L; albumin: 2.8 ± 0.9 g/dl; total bilirubin: 1 ± 0.7 mg/dl).

#### Hepatocytes transplanted into lymph nodes demonstrate long-term survival

To confirm that transplanted hepatocytes were still present in mesenteric lymph nodes at later time points, two of the animals were co-transduced with a lentiviral vector carrying the *Fah* transgene and a second lentiviral vector carrying the *Nis* reporter gene. We performed PET/CT imaging of these two treated pigs to monitor for the expansion of NIS-positive hepatocytes in the mesentery and other tissues. One hour prior to imaging, animals received 10 mCi (370 MBq) of [^18^F]TFB. Pigs 265 and 268 were scanned at 104 and 203 days and 177 and 203 days respectively post-transplantation. Both animals showed NIS-positivity in mesenteric lymph nodes (15.7 and 39.0 SUV and 92.6 and 32.0 SUV, respectively; Fig 3A, Videos S2-5). Despite greater than 50% repopulation of the liver at necropsy with FAH-positive hepatocytes, minimal NIS activity was seen in the liver.

**Figure 3.**
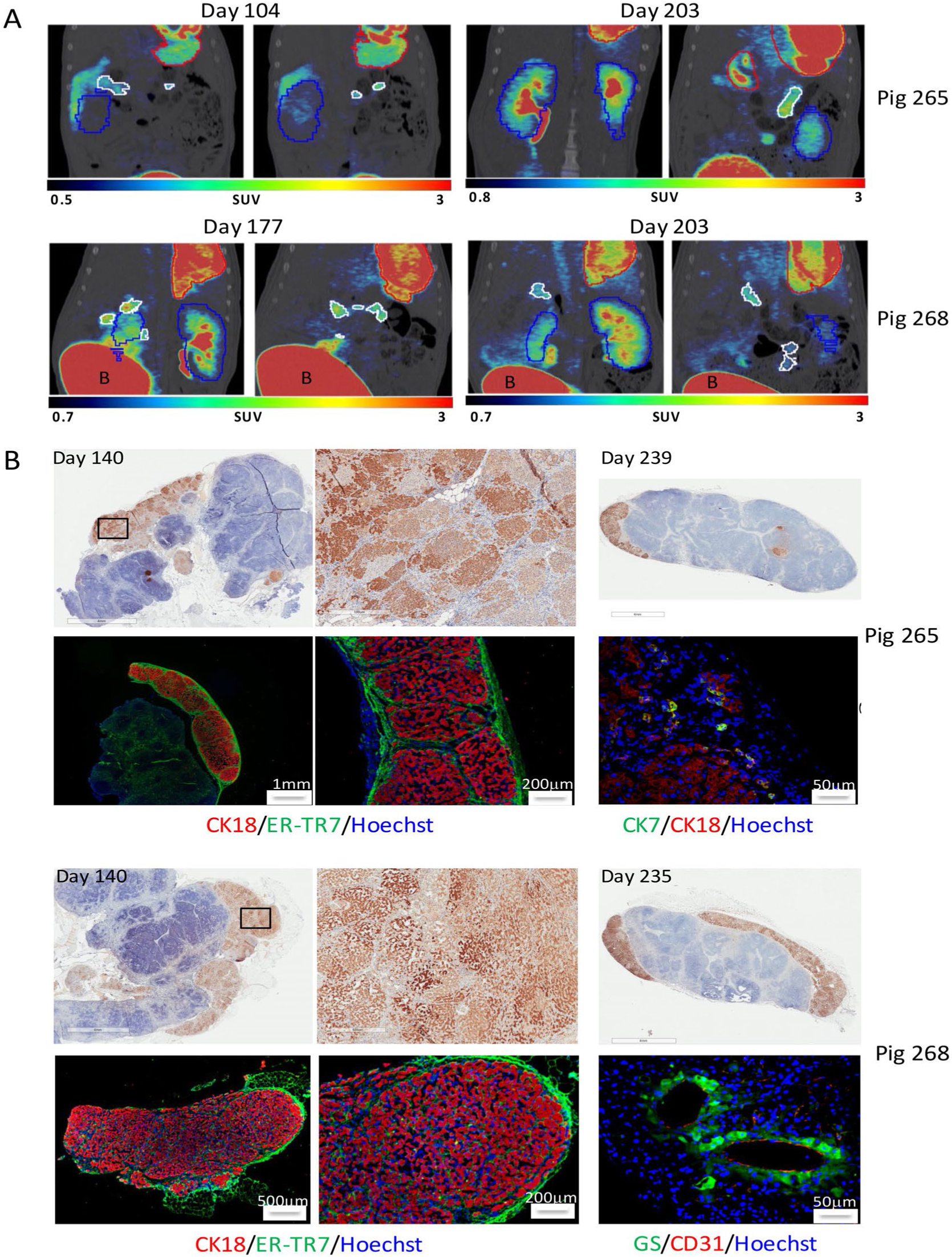
Long-term presence of ectopic liver tissue in mesenteric lymph nodes. (A) PET-CT images of NIS-labeled hepatocytes at 104, 177, and 203 days post-transplantation into mesenteric lymph nodes. Representative coronal images showing persistent engraftment of NIS-labeled hepatocytes in the mesenteric lymph nodes in pigs 265 and 268. Mesenteric lymph nodes are outlined in white; stomach is outlined in red and kidney in blue; bladder is labeled as “B”. (B) Histological confirmation of ectopic hepatocyte presence in mesenteric lymph nodes at 140 and 235-239 days post-transplantation. Pig 265: IHC showing presence of FAH-positive hepatocytes at days 140 (top left, middle) and 239 (top right) post-transplantation. Immunofluorescence at day 239 showing cytokeratin-18 (CK18, red) positive hepatocytes and reticular fibroblasts (ER-TR7, green) on Hoechst background (bottom left, middle), and cytokeratin-7 (CK7, green) positive bile duct cells (bottom right). Pig 268: IHC showing presence of FAH-positive hepatocytes at days 140 (top left, middle) and 235 (top right) post-transplantation. Immunofluorescence at day 235 showing cytokeratin-18 (CK18, red) positive hepatocytes and reticular fibroblasts (ER-TR7, green) on Hoechst background (bottom left, middle), and glutamine synthetase (GS, green) positive hepatocytes around CD31-positive endothelial cells (red) forming possible central vein (bottom right).

To correlate NIS-positivity in mesenteric lymph nodes with histological findings, previously targeted lymph nodes were biopsied in two animals at day 140 post-transplantation. All lymph nodes samples were taken from the root of the mesentery and were grossly positive for hepatic tissue and confirmed by IHC staining for FAH expression, which would be unique to *ex vivo*-corrected hepatocytes due to the use of the *FAH^-/-^* pig and transgenic expression controlled by the liver-specific TBG promoter (Fig 3B). Transplanted hepatocytes were more commonly found in a subcapsular location within the lymph node. All experimental animals were euthanized at 212 to 241 days post-transplantation. Histology of the collected mesenteric lymph nodes showed sustained presence of FAH-positive hepatocytes (Fig 3B, right panels), which was observed in four out of five animals. The two animals that had been biopsied previously still had FAH-positive hepatocyte engraftment in lymph nodes at necropsy confirmed by FAH and cytokeratin 18 IHC staining (Fig 3B, left and middle panels). CD31-positive endothelial cells were found within areas of hepatocyte engraftment in positive tissues (Fig 3B, Pig 268 bottom right panel), suggesting that significant blood vessel remodeling took place after transplantation to sustain long-term engraftment and hepatic growth. Positivity for glutamine synthetase, an enzyme exclusively expressed in pericentral hepatocytes in mammals,(23) was found in hepatocytes surrounding CD31 positive neo-vessels, suggesting neo-formation of a central vein and, importantly, proper physiological zoning of the ectopic liver. Reproduction of hepatic microarchitecture within transplanted lymph nodes was further supported by the presence of hepatic lobules similar to those observed in native pig livers, and cytokeratin 7-positive bile duct cells within the areas of engraftment (Fig 3B, Pig 265 bottom right panel).

### Hepatocytes transplanted into lymph nodes migrate to the liver

Migration of transplanted hepatocytes to the liver was confirmed by FAH IHC of liver tissue demonstrating multiple FAH-positive nodules within the livers of all five animals, covering 67 to 100% (84.7 ± 7.0) of the total liver area at the time of necropsy. A negative correlation was found between percent liver repopulation with FAH-positive hepatocytes and sustained presence of FAH-positive areas of hepatocyte engraftment within lymph nodes; one pig with complete liver repopulation with FAH-positive hepatocytes showed little to no hepatocyte presence remaining within the lymph nodes at the terminal time point, while pigs with only partial liver repopulation with FAH-positive hepatocytes demonstrated a more robust hepatocyte presence within the lymph nodes at 235-239 days post-transplantation (Fig S2). As expected, FAH-negative areas of the liver showed marked hepatocellular damage and fibrosis due to prolonged NTBC withdrawal. However, the two fully FAH-positive livers demonstrated healthy, normal-looking tissue with minimal residual fibrosis, suggesting that the hepatic insult that occurs during NTBC cycling is reversible with time, as FAH-positive hepatocytes expand to repopulate the liver (Fig 4).

**Figure 4.**
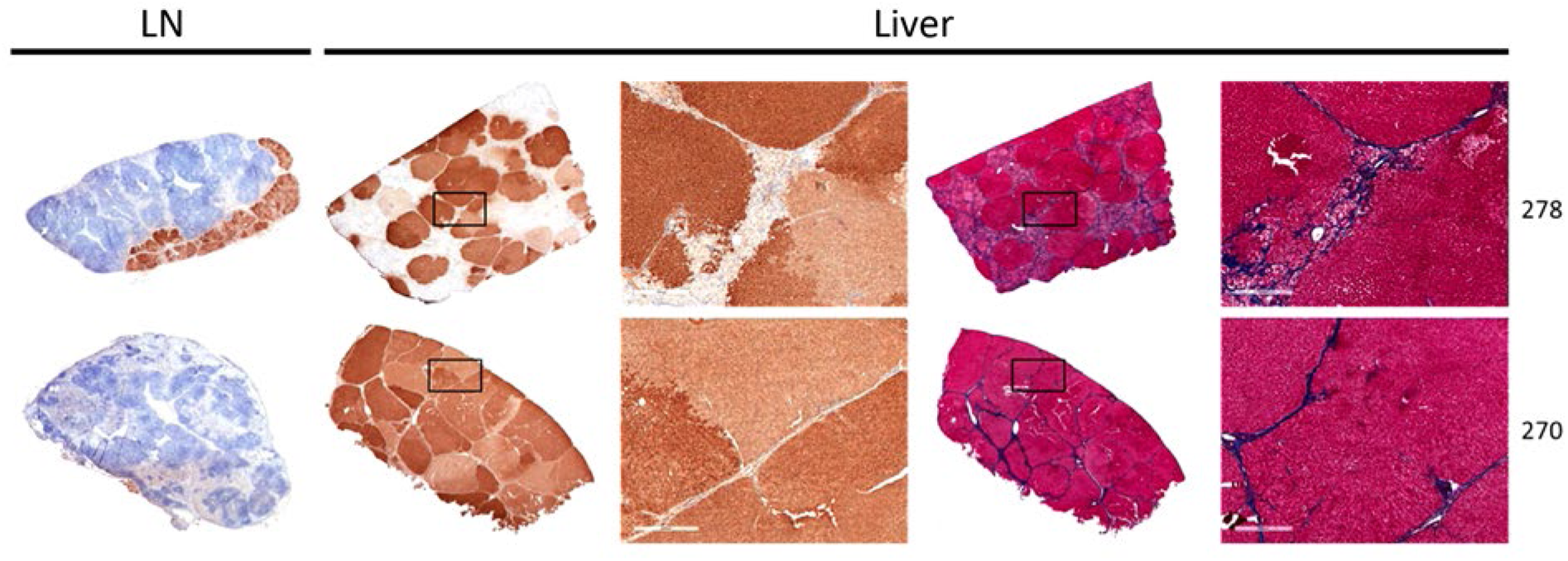
Liver repopulation with FAH-positive hepatocytes. Pig 278: IHC showing presence of FAH-positive hepatocytes in mesenteric lymph node (left) and liver (center) at day 239 post-transplantation; Trichrome (right) showing hepatocellular damage and fibrosis in FAH-negative areas of the liver at day 239 post-transplantation. Pig 270: IHC showing minimal presence of FAH-positive hepatocytes in mesenteric lymph node (left) and complete liver repopulation with FAH-positive cells (center) at day 241 post-transplantation; Trichrome (right) showing resolution of fibrosis with minimal residual scarring in FAH-repopulated livers at day 241 post-transplantation.

In order to characterize any differences between the hepatocytes present in the liver and those present in the lymph nodes, we performed next generation sequencing and bioinformatics analysis of both cell populations. Mapping statistics are provided in Table S1. We found no significant differences in general lentiviral integration profile between these two cell populations. In both cases, integration occurred more often in coding regions than in non-coding regions of the genome, with exons being preferred over introns (Fig 5A). Therefore, integration was favored in chromosomes with higher gene densities (Fig S3B). In both cell populations, lentiviral integration was rare in CpG-rich islands and was clearly favored downstream of transcription start sites (Fig 5B and C), suggesting minimal tumorigenicity potential from oncogene activation. Furthermore, there was no preference for integration in tumor-associated genes in either cell population (Fig 5D).

**Figure 5.**
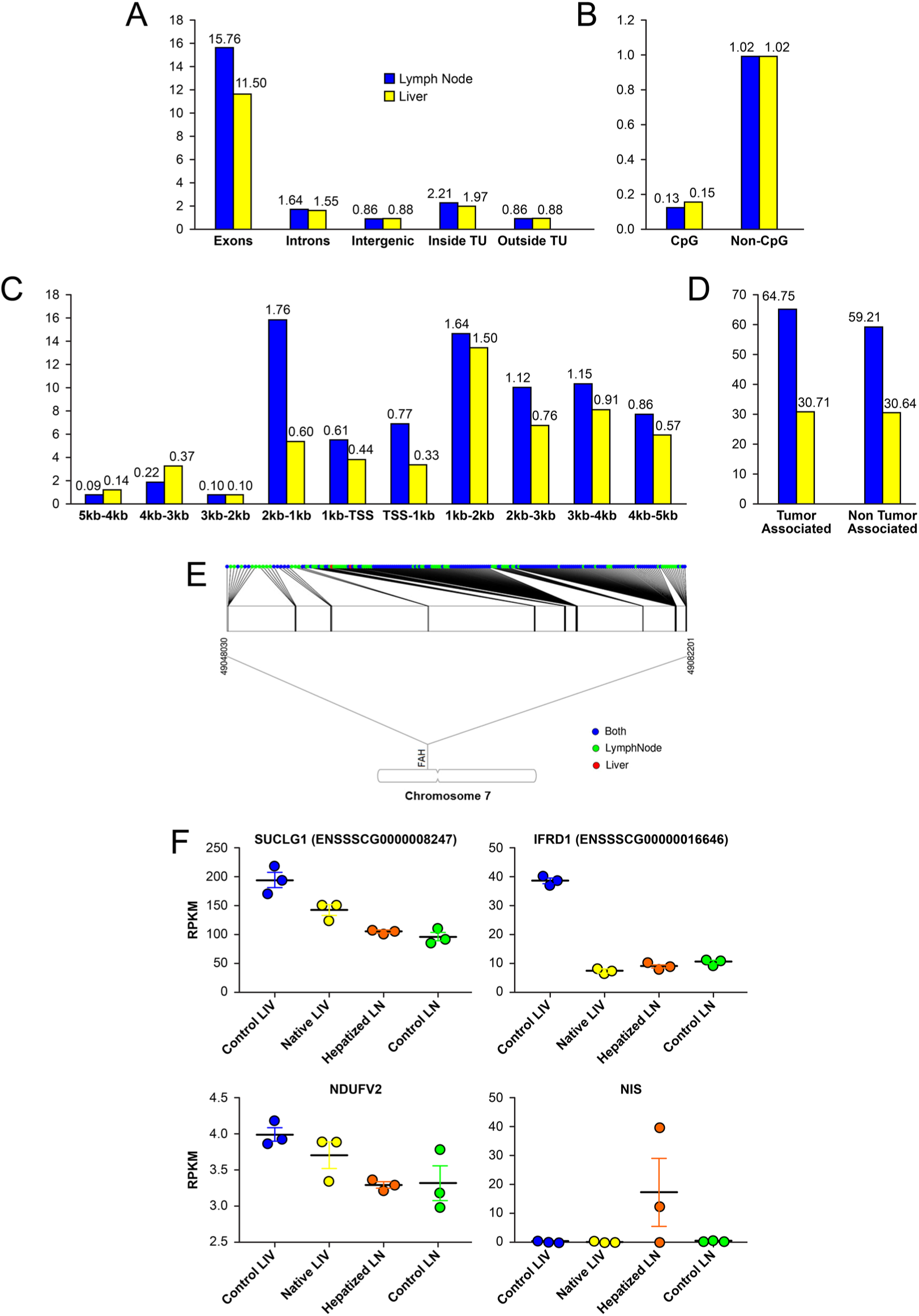
Lentiviral integration profile in lymph node vs. liver hepatocytes. (A) Relative integration frequency in exons, introns, intergenic regions, inside transcription units (TU), and outside transcriptions units for transplanted hepatocytes that remained in lymph nodes and transplanted hepatocytes that migrated to the liver. (B) Relative integration frequency in CpG rich promoter regions vs. other areas of the genome for transplanted hepatocytes that remained in lymph nodes and transplanted hepatocytes that migrated to the liver. (C) Relative integration frequency based on distance upstream and downstream of transcription start sites for transplanted hepatocytes that remained in lymph nodes and transplanted hepatocytes that migrated to the liver. (D) Relative integration frequency in tumor-associated genes vs. other areas of the genome for transplanted hepatocytes that remained in lymph nodes and transplanted hepatocytes that migrated to the liver. (E) Lenti integration into the *fah* locus, 99% of the clones found in the liver were clones of cells that first engrafted in the lymph nodes and then migrated to the liver. (F) Scatter dot plot graphs showing RPKM values for liver-enriched lenti-disrupted genes and the reporter gene (NIS) in control liver, native liver, hepatized lymph node, and control lymph node. Data are mean ± SEM.

The genes with the highest integration frequency in both groups of cells are presented in Table 2, and it is here that differences between the two cell populations were found. *Fah* was the gene with the highest number of distinct integration sites in both groups, although it was only in the top 10 integrated genes in the lymph node population. Interestingly, all but two of the 135 unique integration sites within *Fah* locus in the liver hepatocyte population were represented in the 206 integration sites present in the lymph node hepatocyte population (Fig 5E), indicating that the transplanted cells first engrafted in the lymph nodes, divided and then, after clonal expansion, migrated to the liver. The top four genes with the highest integration frequency in the liver population were also within the top 20 genes in the lymph node population, and were all approximately 5-fold enriched in the liver population when evaluated by the total number of reads present for each, suggesting unbiased expansion and enrichment of a subpopulation of lymph node hepatocytes within the liver. These genes were: *Mir9799*, microRNA; *Ndufv2*, NADH dehydrogenase ubiquinone flavoprotein 2; *Ifrd1*, interferon-related developmental regulator 1; and *Suclg1*, succinyl-CoA ligase (GDP-forming), alpha subunit. In native and ectopic liver samples obtained from our experimental animals after necropsy, expression of *Suclg1, Ndufv2*, and *Ifrd1* was indeed disrupted compared to control liver levels, possibly due to the insertion of the LV construct in at least one allele in some of the corrected hepatocytes (Fig 5F). As a control, NIS expression was evaluated and only detectable in 2 of the 3 samples of hepatized lymph nodes analyzed. The fifth most prevalent integration site in the hepatocytes that migrated to the liver was *Cenpp*, centromere protein P, which was enriched over 78-fold compared to the lymph node population and could be an indication that this disruption caused a favorable migration, engraftment, and/or expansion profile for these clones specifically.

**Table 1.**
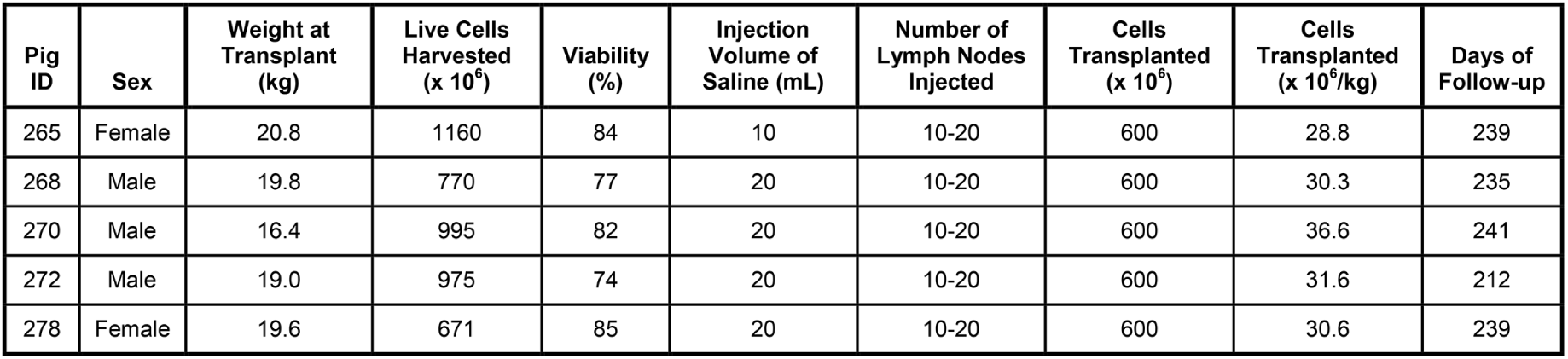
Summary of cell transplantations

| Pig ID | Sex | Weight at Transplant (kg) | Live Cells Harvested (x 10<sup>6</sup>) | Viability (%) | Injection Volume of Saline (mL) | Number of Lymph Nodes Injected | Cells Transplanted (x 10<sup>6</sup>) | Cells Transplanted (x 10<sup>6</sup>/kg) | Days of Follow-up |
| --- | --- | --- | --- | --- | --- | --- | --- | --- | --- |
| 265 | Female | 20.8 | 1160 | 84 | 10 | 10-20 | 600 | 28.8 | 239 |
| 268 | Male | 19.8 | 770 | 77 | 20 | 10-20 | 600 | 30.3 | 235 |
| 270 | Male | 16.4 | 995 | 82 | 20 | 10-20 | 600 | 36.6 | 241 |
| 272 | Male | 19.0 | 975 | 74 | 20 | 10-20 | 600 | 31.6 | 212 |
| 278 | Female | 19.6 | 671 | 85 | 20 | 10-20 | 600 | 30.6 | 239 |

**Table 2.** Genes with most frequent LV-*Fah* integration. Enrichment was calculated as the percent of reads in the liver population divided by the percent of reads in the lymph node population of hepatocytes for specific gene.

| Lymph node Hepatocytes |  |  |  |  | Liver Hepatocytes |  |  |  |  |  |
| --- | --- | --- | --- | --- | --- | --- | --- | --- | --- | --- |
| Top Genes | Number of Integration Sites | Percent of Integration Sites | Number of Reads | Percent of Reads | Top Genes | Number of Integration Sites | Percent of Integration Sites | Number of Reads | Percent of Reads | Enrichment |
| GLRX2 | 26 | 0.40 | 338,309 | 25.57 | MIR9799 | 18 | 0.41 | 5,479,826 | 56.29 | 5.00 |
| MIR9799 | 14 | 0.21 | 148,852 | 11.24 | NDUFV2 | 53 | 1.20 | 1,669,810 | 17.15 | 5.76 |
| LNPK | 53 | 0.81 | 146,239 | 11.05 | IFRD1 | 45 | 1.02 | 952,259 | 9.79 | 5.81 |
| PDCD10 | 81 | 1.24 | 129,982 | 9.82 | SUCLG1 | 24 | 0.54 | 707,921 | 7.27 | 5.06 |
| BEX1 | 51 | 0.78 | 41,620 | 3.15 | CENPP | 77 | 1.75 | 328,807 | 3.38 | 78.69 |
| NDUFV2 | 25 | 0.38 | 39,429 | 2.98 | CPX2 | 36 | 0.82 | 78,896 | 0.81 | 5.63 |
| IDO1 | 5 | 0.08 | 30,360 | 2.29 | CTDSP2 | 45 | 1.02 | 41,364 | 0.42 | 13.71 |
| FAH | 206 | 3.15 | 28,545 | 2.16 | RAB28 | 19 | 0.43 | 23,253 | 0.24 | 6.03 |
| IFRD1 | 31 | 0.47 | 22,270 | 1.68 | QKI | 7 | 0.16 | 16,777 | 0.17 | 5.44 |
| BMP5 | 35 | 0.54 | 19,409 | 1.47 | GSTO1 | 21 | 0.48 | 14,074 | 0.14 | 7.22 |
| MTRF1 | 8 | 0.99 | 19,077 | 1.44 | ENY2 | 30 | 0.68 | 12,470 | 0.13 | 3.97 |
| SUCLG1 | 53 | 0.15 | 19,033 | 1.44 | PRMT6 | 26 | 0.59 | 12,390 | 0.13 | 0.25 |
| ADAMTS1 | 5 | 0.26 | 17,754 | 1.34 | C1QTNF3 | 49 | 1.11 | 11,544 | 0.12 | 1.44 |
| HSD17B3 | 32 | 0.54 | 16,484 | 1.25 | RGS5 | 13 | 0.30 | 11,377 | 0.12 | 2.02 |
| SLC9A2 | 15 | 0.12 | 14,970 | 1.13 | MTRF1 | 36 | 0.82 | 10,972 | 0.11 | 0.08 |
| ACVR2A | 11 | 0.81 | 13,296 | 1.00 | SNCAIP | 5 | 0.11 | 10,252 | 0.11 | 0.81 |
| KCNJ2 | 21 | 0.08 | 12,294 | 0.93 | CTSV | 5 | 0.11 | 9,865 | 0.10 | 5.34 |
| CLDN22 | 8 | 0.49 | 12,073 | 0.91 | FAH | 135 | 3.06 | 9,650 | 0.10 | 0.05 |

To determine whether hepatocytes transplanted in lymph nodes generated a tissue resembling normal liver, transcriptome profiles of untransplanted and hepatized lymph nodes, as well as native (engrafted) and control (isolated from a healthy, untransplanted pig) livers were compared. Twelve thousand genes were identified as differentially expressed (DE) and contrasting control lymph node to liver tissues including hepatized lymph nodes (Figure 6A). T-distributed Stochastic Neighbor Embedding (t-SNE) showed a strong tendency of hepatized lymph node transcripts to cluster with liver tissues samples, whereas control lymph nodes were distinct from the other three groups (Figure 6B). These data suggest that a significant proportion of the lymph node transcriptome acquires a liver-like signature after hepatocyte transplantation. To estimate the similarity of hepatized lymph nodes with control liver, we first focused on 24 genes, which were previously described to be liver-specific (LiGEP, Table S2 and Figure S4). We then included 20 additional genes in our analysis, which can be classified as cytochrome P450 genes, other enzymes, plasma proteins, transporters, surface molecules and cytokines, and transcription factors (Table S2). Cluster heat map showing the RNAseq expression levels of all 44 genes is shown in Figure 6C. Evaluation of albumin expression demonstrates adoption of a liver phenotype by hepatized lymph nodes relative to control (Fig 6D, left) while analysis of *FAH* shows transgenic expression in hepatized lymph nodes as well as elevated transgenic expression in repopulated native liver (under TBG promoter) relative to control liver (Fig 6D, right). Expression levels of all other genes are shown in Fig S4A (LiGEP) and S4B (additional liver-specific genes).

**Figure 6.**
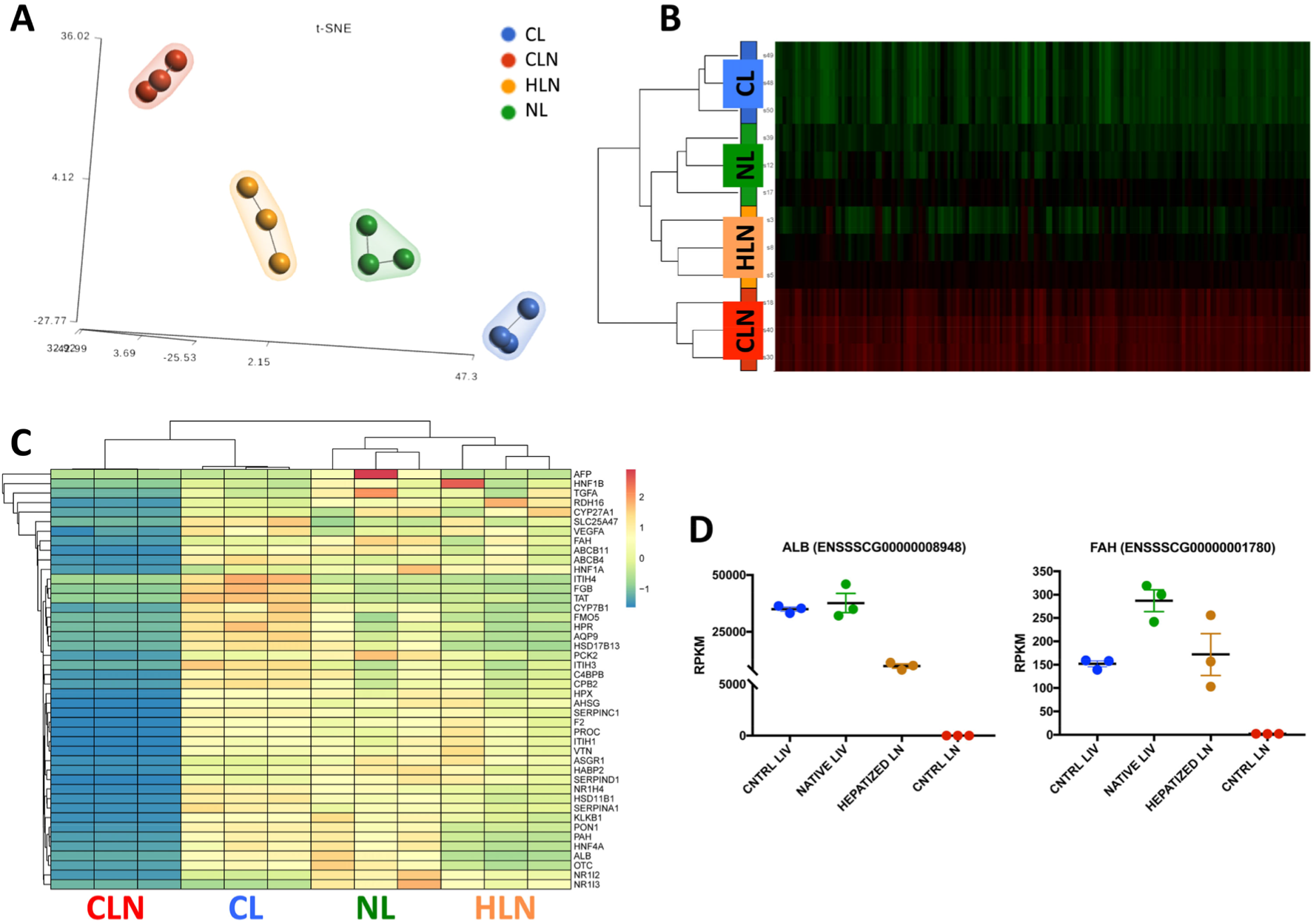
Transcriptional identity of hepatized lymph nodes. (A) t-Distributed stochastic neighbor embedding (t-SNE) 2D plot of RNA-seq data color-coded by tissue sample showing evidence of hepatized lymph node’s (HLN) tendency to cluster with liver samples. CL, control liver; CLN, control lymph node; NL, native liver. (B) Heat map showing the normalized RNA-Seq expression levels (RPKM) of differentially expressed genes (log2 |FC| ≥2, FDR ≥ 0.05, n = 3) across the four conditions. Red and green color intensity indicate gene upregulation and downregulation, respectively, RPKM; Reads Per Kilobase Million. (C) Heat map showing the normalized RNA-Seq expression levels (RPKM) of liver-specific differentially expressed genes (log2 |FC| ≥2, FDR ≥0.05, n = 3) across the four conditions. Spectral colors were used with blue indicating low expression values, yellow indicating intermediately expressed genes, and red representing highly expressed genes. (D) RPKM values for *albumin* (ALB) and *Fah* in control liver, native liver, hepatized lymph node, and control lymph node. Data are mean ± SEM.

## Discussion

In many human liver diseases, orthotopic hepatocyte transplantation and engraftment is often limited by the presence of significant liver injury in the form of inflammation, fibrosis and cirrhosis,(11) underscoring the importance of evaluating alternative transplantation sites. Ectopic hepatocyte transplantation has been attempted in several preclinical and clinical studies.(19) Of the alternative sites tested, only spleen and peritoneal cavity have been able to accommodate a clinically significant number of cells.(24) Intrasplenic injection of hepatocytes improves liver function and prolongs survival in a mouse model of liver failure.(25) Although transsplenic access to the portal vein in human patients has been done (26), this may introduce hemorrhagic risk that is decreased with lymph node transplantation. Intraperitoneal transplantation of hepatocytes has been attempted in humans with acute liver failure, but transplanted cells are not able to survive in this environment for extended periods of time.(27)

Lymph nodes are a highly favorable site for ectopic cell transplantation.(19) Not only does their plasticity and highly complex vascular network provide a healthy milieu for cell engraftment and expansion, as demonstrated by both their natural interactions with lymphocytes and their pathological interactions with metastatic tumor cells, but their accessibility makes it possible to perform minimally invasive cell delivery in the clinical setting. This could prove especially important in patients that are higher risk surgical candidates. There is ample precedent in the clinic for ultrasound-guided lymph node identification and targeting. Fine needle aspiration biopsies, for example, are now a routine diagnostic test. Multiple clinical studies have also made use of lymph nodes as sites for injection of immunotherapeutic agents, with patients rating the procedure comparable to a subcutaneous injection and less painful than venous puncture.(28) Furthermore, cell transplantation into lymph nodes does not appear to affect their function: no complications were noted in our study or in previous studies.(20) (21)

All five animals treated through this method demonstrated NTBC independence within three to five months of treatment, as well as complete normalization of tyrosine levels and liver function tests. This timeframe to metabolic correction is comparable to that of our previous study using similar numbers of cells and portal vein delivery into the native FAH-deficient liver.(10) It is worth noting that in our previous study, animals were also maintained on the protective drug NTBC until the time of transplantation, which means that their livers did not have a significant amount of inflammation and fibrosis when they received the corrected hepatocytes. Orthotopic cell transplantation has shown low rates of engraftment, especially when the liver is compromised by inflammation and fibrosis, as is seen in many liver diseases.(29)

It has been previously shown that hepatocyte transplantation into lymph nodes is able to generate enough ectopic liver mass to correct HT1 and rescue mice from acute and chronic liver disease.(21) In our study, we show a similar result in the large animal model of HT1, suggesting that hepatocyte transplantation into lymph nodes is scalable and may therefore be a clinically relevant technique that permits the creation of sufficient ectopic liver mass to significantly impact liver function. There are important differences between these two preclinical studies. Syngeneic and allogeneic hepatocytes were used in the HT1 mouse study by Komori *et al*, and the HT1 mouse does not exhibit the full extent of liver injury seen in humans, including fibrosis and cirrhosis.(21) The current study was conducted in the HT1 pigs initially supported on NTBC, where liver disease could be delayed until the harvesting of autologous cells was complete. In this study we take advantage of the porcine HT-1 model of liver failure because 1-2 weeks after the withdrawal of NTBC we begin to see rapid and severe liver injury that includes all the same features we see in human liver failure including, inflammation, fibrosis and cirrhosis. This study design allows us to isolate the single variable related to ectopic transplantation of hepatocytes in LNs in the setting of liver failure while avoiding potential confounding variables related to immune suppression or the use of primary hepatocyte alternatives. Now that we have demonstrated the feasibility of targeting LNs for hepatocyte transplantation, subsequent evaluations should be performed in untreated HT1 pigs or other large animal models of liver disease using primary hepatocyte alternatives or allotransplantation and immune suppression.

The fact that our HT1 model, which develops all the acute and chronic liver failure features seen in many other liver diseases, was completely weaned of NTBC requirement and metabolically normalized after the ectopic transplantation procedure indicates the therapeutic potential of this approach. Even though the goal of this study was not to evaluate directly the use of ectopic LN hepatocyte transplantation for the treatment of any specific genetic liver disease, there are some potential applications that should be discussed regarding these diseases. Some optimizations in initial engraftment rates or other modifications (i.e., exogenously applied selective pressure or repeat administrations of corrected cells) would be necessary for diseases that do not inherently propagate healthy cells, such as phenylketonuria or Wilson’s disease. However, it is the liver damage itself through the hepatectomy and not something specific to HT1 that provides the stimulus for initial successful engraftment. Additionally, unlike HT1, most liver diseases do not require complete repopulation of the diseased liver, and administration could be catered to effect to see a phenotypic change.

Autologous hepatocyte transplantation could be directly translated into human patients where their condition does not preclude a partial hepatectomy, thereby avoiding immune concerns related to the use of allogeneic cells. However, healthy allogeneic hepatocytes could be transplanted in cases where partial hepatectomy is contraindicated, such as in acute liver failure or severe cirrhosis, which would require additional support via established immune suppression protocols. Possible future sources of hepatocytes include universal donor hepatocytes, IPS-derived hepatocytes, and HLA-matched farmed hepatocytes (30-32). In our study hepatocytes were transplanted into pig mesenteric lymph nodes using an open technique due to the paucity of suitable peripheral lymph nodes in the neonatal pig, but the same procedure could foreseeably be performed into central or peripheral lymph nodes in humans via a percutaneous or endoscopic ultrasound-guided technique, as described previously.(28)

In our model, we found that although hepatocytes demonstrated long-term survival within lymph nodes, forming appropriate liver architecture including bile ducts, some also subsequently engrafted in the liver after a period of engraftment and expansion in the LNs. This was suggested by bioinformatics data and confirmed by histology. This phenomenon was very limited in previous mouse studies(21) and could be related to the microanatomy and direction of flow in pig lymph nodes, which differ from those of other mammals. Their microarchitecture is inverted, with the germinal centers being located internal to the medulla, which has led authors to suggest that lymphocytes are transported back into a capillary system after passing through a lymph node, as opposed to the lymphatic system as in other animals.(33, 34) Indeed, during the injection procedure it was possible to observe injected fluid communicating from the lymph node into the capillary bed.

The fate of the engrafted hepatocytes into the lymph nodes was assessed by the evaluation of lentiviral vector integration into prevalent site, such as the *Fah* locus. Of 135 unique integration sites in the *Fah* locus seen in the hepatocytes engrafted in the liver, only 1 was not among the 206 unique integration sites observed in the hepatocytes in lymph nodes. This result substantiates the conclusion that although cells migrate from the lymph nodes to the liver after transplantation they do so primarily after a period of engraftment in the lymph nodes. If the hepatocytes had migrated immediately to the liver after transplantation we would have expected a more disparate integration profile between the two populations. Further experiments conducted on a different large animal model with a similar lymphatic anatomy to humans would be required to judge whether the level of hepatocyte migration from mesenteric lymph nodes to liver remains the same. If so, central lymph node injection could potentially be considered a safer alternative to portal vein injection for hepatocyte delivery into the liver, since portal vein infusion of cells often leads to thrombotic complications and requires systemic anticoagulation.(35)

Based on these results, we also analyzed in detail the overall integration profile of our lentiviral vector in both hepatocytes that remained in the lymph nodes and hepatocytes that migrated to the liver. In both cell populations, the lentiviral vector showed a benign integration profile, with no preference for integration in promoter regions or tumor-associated genes. Analysis of the genes with the highest integration frequency in both populations suggested that hepatocytes that migrated to and expanded in the liver were a subpopulation of hepatocytes that remained in the lymph nodes, although discerning any functional impact of disrupting the integrated genes was beyond the scope of the current study. Interestingly, prevalence of the top four integrated genes in the liver population was approximately five-fold higher than in the lymph node population, a linearity possibly supported by a clonal founder with all four integrations or up to 4 unique clonal founders with identical (unbiased) expansion after migration and liver engraftment. Of note, *Fah* was within the top twenty genes with the highest integration frequency and was the gene with the highest number of distinct integration sites in both groups. This suggests that homology may guide integration to some extent, a finding that provokes further development to possibly improve safety and predictability of lentiviral genomic disruption.

We also examined relative changes in gene expression, finding no significant differences in general transcriptomes between engrafted and control lymph nodes, as well as engrafted and control liver, aside from specific variations induced by transgene expression (i.e. *FAH*) or lymph node adoption of a liver phenotype (i.e., albumin expression). This implies that transplantation of hepatocytes in lymph nodes and *ex vivo* lentiviral gene therapy does not significantly alter endogenous gene expression within the corrected cells. Together with the integration profiling data (discussed above), this advocates for the safety of the development of an ectopic liver in lymph nodes and the *ex vivo* lentiviral gene therapy approach. Given the clinical success of *ex vivo* lentiviral gene therapy in cancer via CAR-T cells (see review by Geyer and Brentjens, 2016) and the over 250 open FDA trials using lentivirus, our data support progression of this paradigm into additional indications. Ultimately, approval of expanded clinical application of *ex vivo* lentiviral gene therapy will be made on a risk:reward basis, and a growing body of literature from our laboratories and others support both safety (reduced risk) and efficacy (increased reward) of this approach.

In summary, we have shown that ectopic transplantation of corrected, autologous hepatocytes into mesenteric lymph nodes is curative in the HT1 pig model of severe liver failure. Cells transplanted into these ectopic sites engraft and expand creating hepatized LNs, aiding the failing liver. Hepatocyte transplantation into lymph nodes is a promising approach to the treatment of multiple liver diseases that holds several important advantages over more traditional cell transplantation techniques, especially in patients with pre-existing native liver damage.

## Materials and Methods

### Study design

This prospective controlled laboratory experiment was designed to address the hypothesis that ectopic transplantation of hepatocytes would result in sufficient functional biomass to support a large animal model of liver failure, specifically the porcine model of hereditary tyrosinemia type I (HTI) during NTBC withdrawal. Five HT1 pigs underwent *ex vivo* gene therapy with two lentiviral vectors (one expressing FAH and the other expressing the NIS reporter gene). The NIS reporter should not affect phenotype but is only intended to allow non-invasive tracking of transplanted cells. The finding that ectopic transplantation of corrected hepatocytes seeded subsequent robust orthotopic engraftment was not anticipated, and the additional hypothesis that the hepatocytes that migrated to liver were a subset of those that were initially engrafted in the lymph nodes (as opposed to a unique population that never engrafted ectopically) was further developed and tested using bioinformatics.

These studies in large animals were performed in a limited number of subjects (n=5) due to resource-intensive nature of the involved surgical and diagnostic procedures in large animals, and associated resource requirements for the husbandry, *in vivo* evaluations, and bioinformatic analyses.

### Animals

All animal procedures were performed in compliance with Mayo Clinic’s Institutional Animal Care and Use Committee regulations and all animals received humane care. For biodistribution experiments, a female wildtype pig was used. For *ex vivo* gene therapy experiments, male and female *Fah*^-/-^ pigs were used. *Fah*^-/-^ pigs were produced in a 50% Large White and 50% Landrace pig as previously described.(36, 37)

### NTBC administration

NTBC mixed in food was administered at a dose of 1 mg/kg/day with a maximum of 25 mg/day. All animals remained on NTBC until the time of transplantation, after which NTBC administration was discontinued to support expansion of the corrected cells. After hepatocyte transplantation, all animals were monitored daily for loss of appetite or any other clinical signs of morbidity. Animals were weighed daily for the first two weeks post-operatively and weekly thereafter. If loss of appetite, weight loss, or any other signs of morbidity occurred, NTBC treatment was reinitiated for seven days. Animals were cycled on and off NTBC in this fashion to stimulate expansion of corrected FAH-positive cells.

### Liver resection and hepatocyte isolation

Eight-week-old (16-21 kg) pigs underwent a laparoscopic partial hepatectomy involving the left lateral lobe under inhaled general anesthesia with 1-3% isoflurane. Resection volumes represented 15 to 20% of the total liver mass. Using an open Hasson Technique, a 12 mm port was placed into the peritoneal cavity and the abdomen insufflated. Using a 5 mm laparoscope (Stryker, Kalamazoo, MI), two additional 5 mm ports were placed under direct visualization. The left lateral segment and its vascular and biliary drainage was isolated and divided using an Endo GIA stapler (Covidien, Dublin, Ireland). The liver section was retrieved using an Endo Catch bag (Covidien, Dublin, Ireland), and adequate hemostasis was ensured prior to port removal and incision closure. This liver section was then perfused *ex vivo* through the segmental portal vein with a two-step perfusion system to isolate hepatocytes as previously described.(38) Number and viability of cells were determined by trypan blue exclusion.

### Hepatocyte radiolabelling

For early biodistribution experiments, hepatocytes were radiolabeled in suspension with ^89^Zr (t_1/2_= 78.4 hr) with synthon ^89^Zr-DBN at 27°C for 45 min in Hank’s Buffered Salt Solution (HBSS) as previously described. (39-41)

### Hepatocyte transduction

Hepatocytes were co-transduced in suspension at a MOI of 20TUs with two third-generation lentiviral vectors carrying the sodium-iodide symporter (*Nis*) reporter gene or the pig *Fah* gene under control of the thyroxine-binding globulin promoter (Fig 1A). The NIS reporter allows for longitudinal non-invasive monitoring of transplanted cells through single-photon emission computed tomography (SPECT) or positron emission tomography (PET).(42) Transduction occurred over the course of 2 h before transplantation using medium and resuspension techniques previously described.(10) Hepatocytes were resuspended in 0.9% sodium chloride (Baxter Healthcare Corporation, Deerfield, IL).

### Hepatocyte transplantation in pigs

All pigs received autologous transplantation of hepatocytes through direct mesenteric lymph node injection (Fig 1B). After partial hepatectomy, animals were kept under general anesthesia until the time of transplantation, approximately 4 h later.. Bowel was exteriorized through the upper midline incision until the root of the mesentery was visible (Fig 1C). Mesenteric lymph nodes were identified and hepatocytes were delivered directly into 10-20 nodes through direct injection with a 25-gauge one-inch needle. Each animal received a total of 6 x 10^8^ hepatocytes in 10-20 mL of saline. Heparinization of the cell solution just prior to injection was performed at 70 U/kg of recipient weight as has been previously described for islet cell transplantation protocols.(43)

### Positron emission tomography–computed tomography (PET-CT) imaging and analysis

Imaging was performed on the high-resolution GE Discovery 690 ADC PET/CT System (GE Healthcare, Chicago, IL) at 6, 54, and 150 h post-transplantation in the ^89^Zr-labeled animal. Noninvasive evaluation of NIS expression was performed by PET/CT at 3, 5, and 6 month post-transplantation in two of the NIS-labeled animals using [^18^F]tetrafluoroborate ([^18^F]TFB) radiotracer.(44) CT was performed at 120 kV and 150 mA, with tube rotation of 0.5 s and pitch of 0.516. PET was performed as a two-bed acquisition with 10 min per bed and 17-slice overlap, resulting in a 27-cm axial field of view. PMOD (version 3.711; PMOD Technologies, Switzerland) was used for image processing, 3D visualization and analysis. The anatomic location of liver and kidneys was identified from the registered datasets. Volumes of interest (VOIs) of mesenchymal lymph nodes were created on PET images, and standardized uptake values (SUVs) for body weight were obtained. Surface rendering was performed using the threshold pixel value for bone, liver and kidneys from the CT dataset and lymph nodes from the PET dataset to localize the lymph nodes with reference to the skeleton.

### Biochemical analysis

Standard serum and plasma analyses were performed. Tyrosine values in plasma were determined using liquid chromatography and tandem mass spectrometry via Mayo Clinic’s internal biochemical PKU test.

### Histopathological and immunohistochemical analyses

For H&E, Masson’s Trichrome and FAH stains, tissue samples were fixed in 10% neutral buffered formalin (Azer Scientific, Morgantown, PA), paraffin embedded and sectioned (5 μm). H&E and Masson’s Trichrome stains were performed by means of standard protocols, while immunohistochemistry for FAH was performed as previously described.(42) Percent of FAH-positive hepatocytes was quantified using a cytoplasmic stain algorithm in Aperio ImageScope. For all other immunohistochemical stains, tissues were fixed in 4% paraformaldehyde for 4 h, infiltrated with 30% sucrose overnight and then embedded in OCT compound prior to sectioning (5-7 μm). Sections were washed with phosphate-buffered saline (PBS) and blocked with 5% bovine serum albumin (BSA) or milk for 30 min. Sections were then incubated with primary antibody for 1 h and secondary antibody for 30 min, and were coverslipped with a glycerol-based mounting media containing Hoechst. Images were captured with an Olympus IX71 inverted microscope. The following primary antibodies were used: ER-TR7 (ab51824, Abcam, Cambridge, MA), Glutamine Synthetase (ab64613, Abcam, Cambridge, MA), CD31 (MCA1746GA, Bio-Rad, Hercules, CA), Cytokeratin 7 (ab9021, Abcam, Cambridge, MA) and Cytokeratin 18 (10830-1-AP, Proteintech, Chicago, IL).

### Amplification, next-generation sequencing (NGS), and genomic DNA mapping of lentiviral integration sites

Genomic DNA was isolated from snap-frozen tissue fragments of hepatized mesenteric lymph nodes and engrafted livers from Animal No. 265 using a Gentra Puregene Tissue Kit (Qiagen, Hilden, Germany). Ligation-mediated PCR (LM-PCR) was used for efficient isolation of integration sites. Restriction enzyme digestions with MseI were performed on genomic DNA samples; the digested DNA samples were then ligated to linkers and treated with ApoI to limit amplification of the internal vector fragment downstream of the 5′ LTR. Samples were amplified by nested PCR and sequenced using the Illumina HiSeq 2500 Next-Generation Sequencing System (San Diego, CA). After PCR amplification, amplified DNA fragments included a viral, a pig and sometimes a linker segment. The presence of the viral segment was used to identify reads that report a viral mediated integration event in the genome. The reads sequenced from these DNA fragments were processed through quality control, trimming, alignment, integration analysis, and annotation steps.

Quality control of the sequenced read pairs was performed using the FASTQC software. The average base quality of the sequenced reads was >Q30 on the Phred scale. Uniform distribution of A,G,T and C nucleotides was seen across the length of the reads without bias for any specific bases. The number of unknown “N” bases was less than 1% across the length of the reads. High sequence duplication (> 80%) was observed, however, this was anticipated due to the nature of the experiment and the amplification of specific library fragments.

We used Picard software’s (http://broadinstitute.github.io/picard/) insert size metrics function to calculate the average fragment length per sample. This metric averaged across all the samples was 158 base pairs (bps) long with an average standard deviation of 29bps. Since 150bp long reads were sequenced, most of the paired read one (R1) and read two (R2) reads overlapped, providing redundant information. R2 reads were therefore not considered in the analysis.

Reads were trimmed to remove the viral sequence in two separate steps. In step 1, the viral sequence was trimmed from the R1 reads using cutadapt(45) with a mismatch rate (e=0.3) from the 5’ end of each read. In step 2, if the linker sequence was present, it was similarly trimmed from the 3’end of the read. Trimmed reads with a length less than 15bps were removed from the rest of the analysis to reduce ambiguous alignments. Untrimmed reads were also removed from the rest of the analysis because they did not contain a viral segment that could be used as evidence of a viral mediated integration.

The remaining R1 reads were aligned to the susScr11 build of the pig reference genome using BWA-MEM in single-end mode.(46) Default BWA-MEM parameters were used. The reads used to identify genomic points of viral integrations had to be uniquely mapped to the genome with a BWA-MEM mapping quality score greater than zero.

An integration point was defined by the position of R1’s first aligned base on the susScr1 genome. Unique integration points were identified across the genome without any constraints on coverage. However, for downstream annotation and analysis, only those integration points with 5 or more supporting reads were used to minimize calling false positive integration points.

Locations of integration points were categorized in: exons, introns, 3 prime UTRs, 5 prime UTRs and intergenic regions using information extracted from the susScr11 refflat file maintained by UCSC’s Genomics Institute. To avoid conflict of feature categorization arising from multiple overlapping gene definitions, only the definition of the longest gene was used to annotate integration points. Additional features where computed including: the distance to the nearest transcription start site (TSS), makeup of CpG-rich regions of the genome (“CpG islands”), and enrichment to tumor-associated genes. This tumor gene list was comprised of 745 clinically validated tumor-associated human genes (from Mayo Clinic internal data). Those genes were related to their pig homologs based on GenBank’s gene names.

### RNA isolation, tissue screening, and whole transcriptome sequencing (RNAseq)

Total RNA was isolated from snap-frozen tissue fragments of hepatized or control (non-transplanted) mesenteric lymph nodes from three experimental animals, and of engrafted livers from the same three experimental animals or control liver from one untreated, control animal using the RNeasy Mini Kit (Qiagen, Valencia, CA) according to manufacturer’s instructions (including the optional on-column DNase digestion). RNA purity and concentration were measured using a NanoDrop 2000/c spectrophotometer (Thermo Fisher Scientific, Pittsburgh, PA). Screening of hepatized lymph nodes was performed by examining *albumin* and *fah* expressions using quantitative reverse transcription PCR (qRT-PCR). Non-transplanted mesenteric lymph nodes and a control liver were used as negative and positive controls, respectively. RNA was retrotranscribed using the iScript Reverse Transcription Supermix (Bio-Rad, Hercules, CA) and *albumin* and *fah* were amplified using the SsoAdvanced™ Universal SYBR® Green Supermix (Bio-Rad, Hercules, CA) on a StepOnePlus Real-Time PCR System (Applied Biosystems, Foster City, CA). Expression of glyceraldehyde-3-phosphate dehydrogenase (GAPDH) was used for normalization of gene expression data. Relative changes in gene expression were calculated using the 2^-ΔΔCT^ method. RNAs from selected samples were shipped on dry ice to Novogene in Sacramento, CA for library preparation and sequencing. All samples passed Novogene internal quality control with RNA integrity number above eight.

RNAseq data analysis was performed using Partek Genomics Suite software version 7.0. Briefly, quality control was measured considering sequence-read lengths and base-coverage, nucleotide contributions and base ambiguities, quality scores as emitted by the base caller, and over-represented sequences. All the samples analyzed passed all the QC parameters and were mapped to the annotated pig reference SScrofa11.1_90+nonchromosomal using STAR2.4.1d index standard settings. Estimation of transcript abundance based on the aligned reads was performed by optimization of an expectation-maximization algorithm and expressed as FPKM (Fragments Per Kilobase Million). Differentially expressed (DE) genes were detected using differential gene expression (GSA) algorithm based on p-value of the best model for the given gene, false discovery rate (FDR), ratio of the expression values between the contrasted groups, fold change (FC) between the groups and least-square means (adjusted to statistical model) of normalized gene counts per group. 12000 genes were identified as DE based on log2 |FC| ≥2 and FDR ≥ 0.05 cut-off. Unsupervised clustering to visualize expression signature was performed using 1-Pearson correlation distance and complete linkage rule and samples were classified using t-SNE (t-distributed stochastic neighbor embedding).

### Statistical analysis

Numerical data are expressed as mean ± SD (standard deviation) or SEM (standard error of the mean). Statistical significance was determined by Welch’s t-test, and established when p ≤ 0.05. Statistical analyses were performed with GraphPad Prism software version 7.

## Supporting information

Sup Vid 1

Sup Vid 2

Sup Vid 3

Sup Vid 4

Sup Vid 5

## Acknowledgements

We thank T. Wyman and L. Filzen for large animal PET imaging support; L. Gross and L. Acosta for histology support; S. Krage and J Pederson for surgical support; L. Hillin for animal support. P. Steiner for graphical support.

## Author contributions

CN, RH, RK, and JL conducted experiments and wrote the manuscript. SN designed experiments. EL designed experiments, interpreted results, and revised the manuscript. BH, MF, AC, KA, RG, CV, BA conducted experiments. ZD developed reagents and conducted experiments. LS, HJ, AB, MP, IG, VL, AB, DO, JK, TD, provided expertise and interpreted data.

## Data and materials availability

The datasets generated and/or analyzed during the current study are available from the corresponding author on reasonable request.

## Supplementary Materials

**Supplementary Figure 1.**
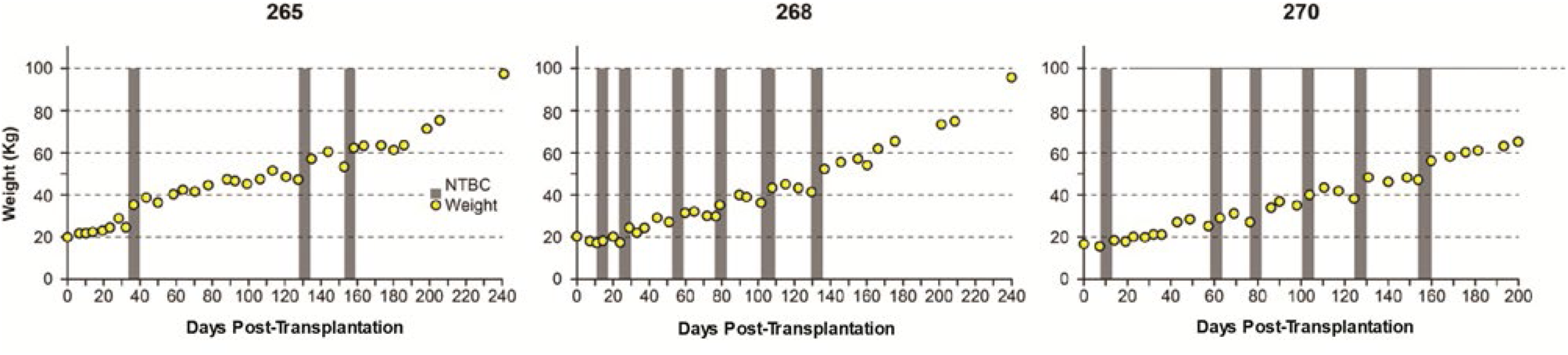
Weight stabilization of pigs 265, 268, and 270. NTBC-independent weight gain was achieved at days 158, 133, and 163, respectively, after three to six cycles on the drug.

**Supplementary Figure 2.**
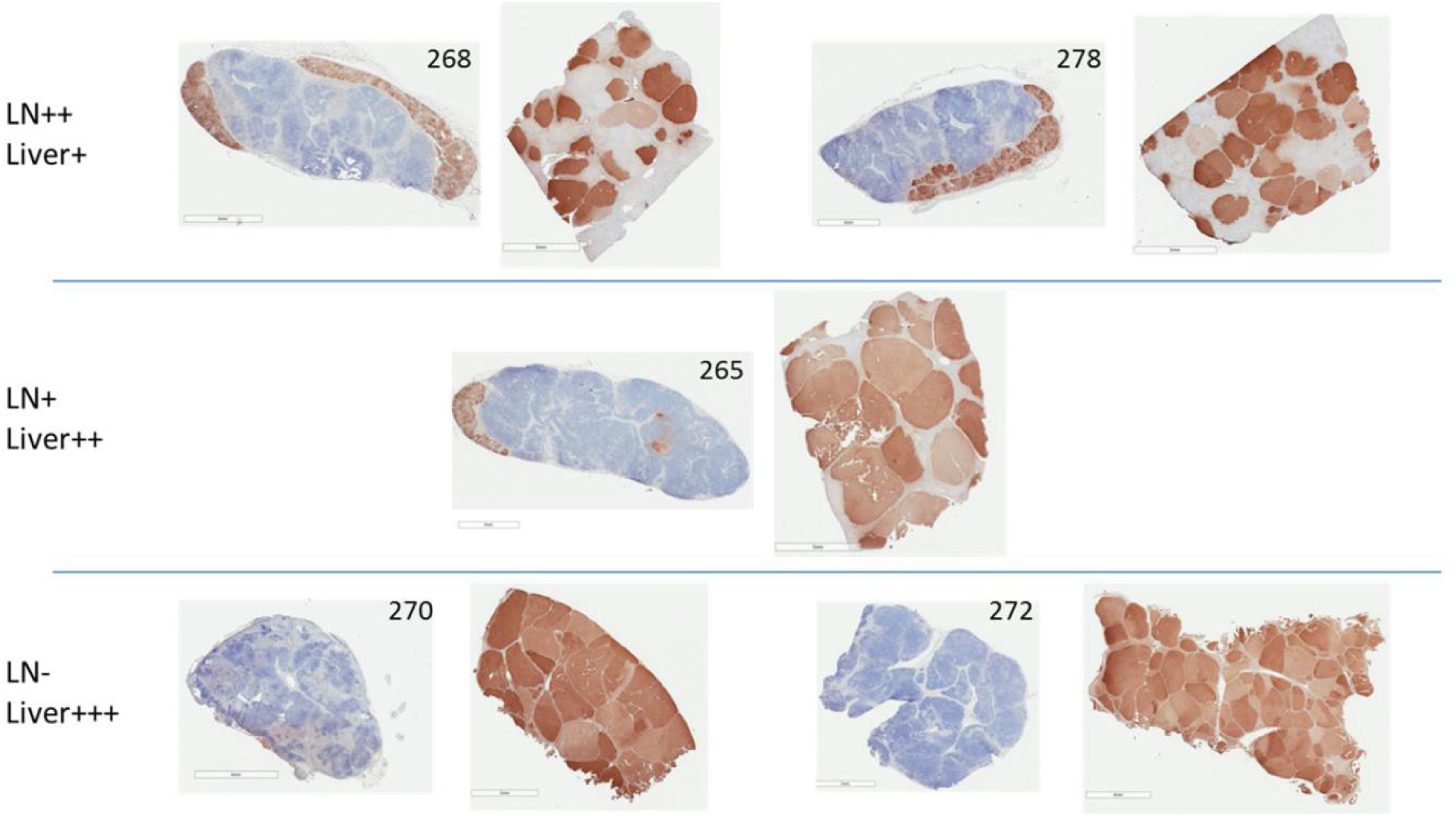
Correlation between presence of FAH-positive hepatocytes in lymph node and liver. Animals with partial liver repopulation with FAH-positive hepatocytes (pigs 268 and 278) still had robust ectopic liver tissue presence in lymph nodes at the time of euthanasia. Animals with complete liver repopulation with FAH-positive hepatocytes (pigs 270 and 272) had little to no presence of ectopic liver tissue in lymph nodes at the time of euthanasia.

**Supplementary Table 1.** Next generation sequencing mapping statistics

| Sample | Total Reads | Mapped Reads (Total) | Integration Points at 1X | Integration Points at 5X |
| --- | --- | --- | --- | --- |
| Lymph node | 115.748.080 | 5,465,009 (7.34 %) | 40.239 | 6.335 |
| Liver | 116.490.845 | 21,126,444 (22.29 %) | 27.987 | 4.270 |

**Supplementary Figure 3.**
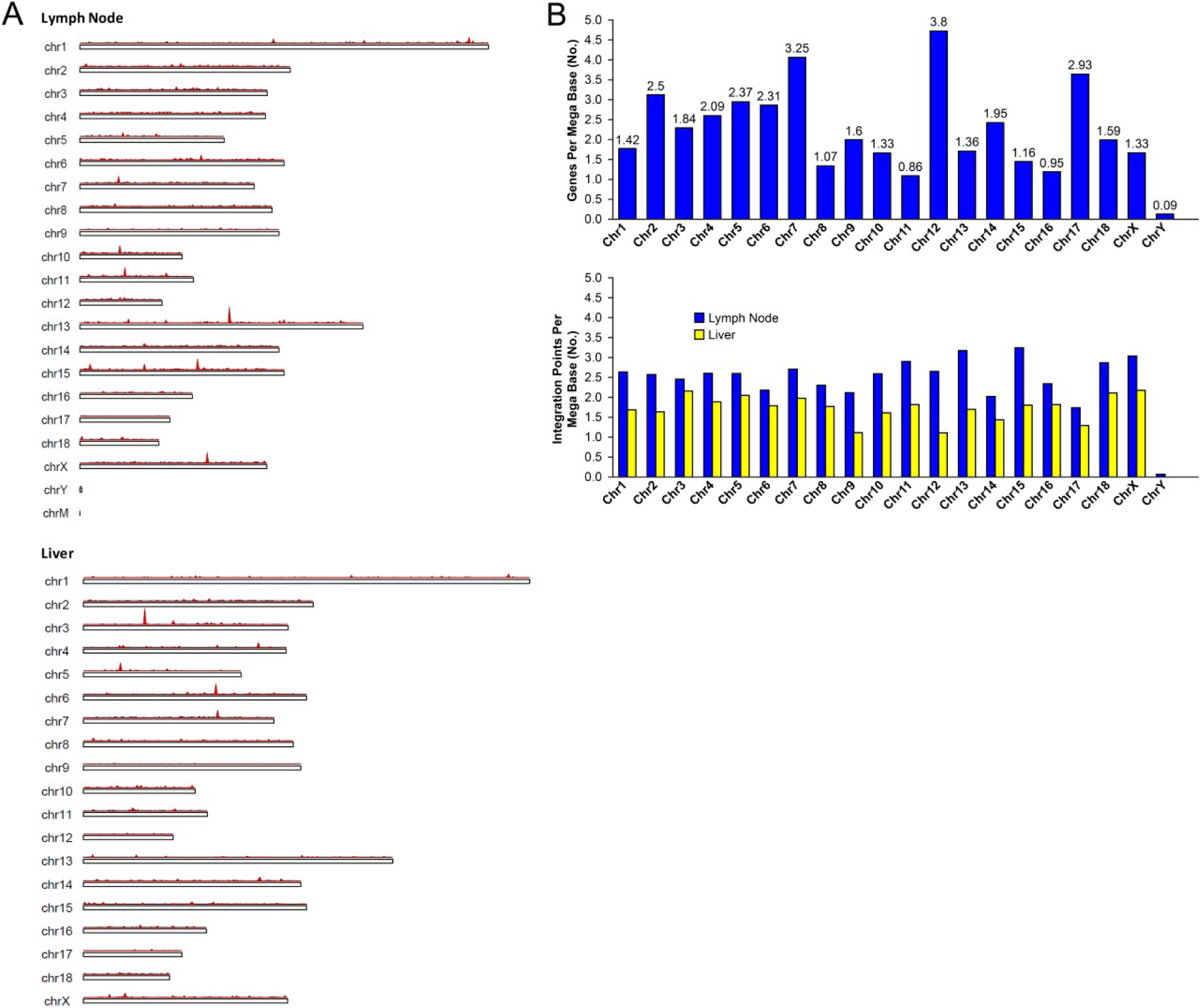
Lentiviral integration profile in lymph node vs. liver hepatocytes. (A) Chromosomal integration map for transplanted hepatocytes that remained in lymph nodes (top) and transplanted hepatocytes that migrated to the liver (bottom). Red vertical bars represent integration points. (B) Relative gene density per chromosome (top) compared to relative integration density per chromosome (bottom).

**Supplementary Table 2.**
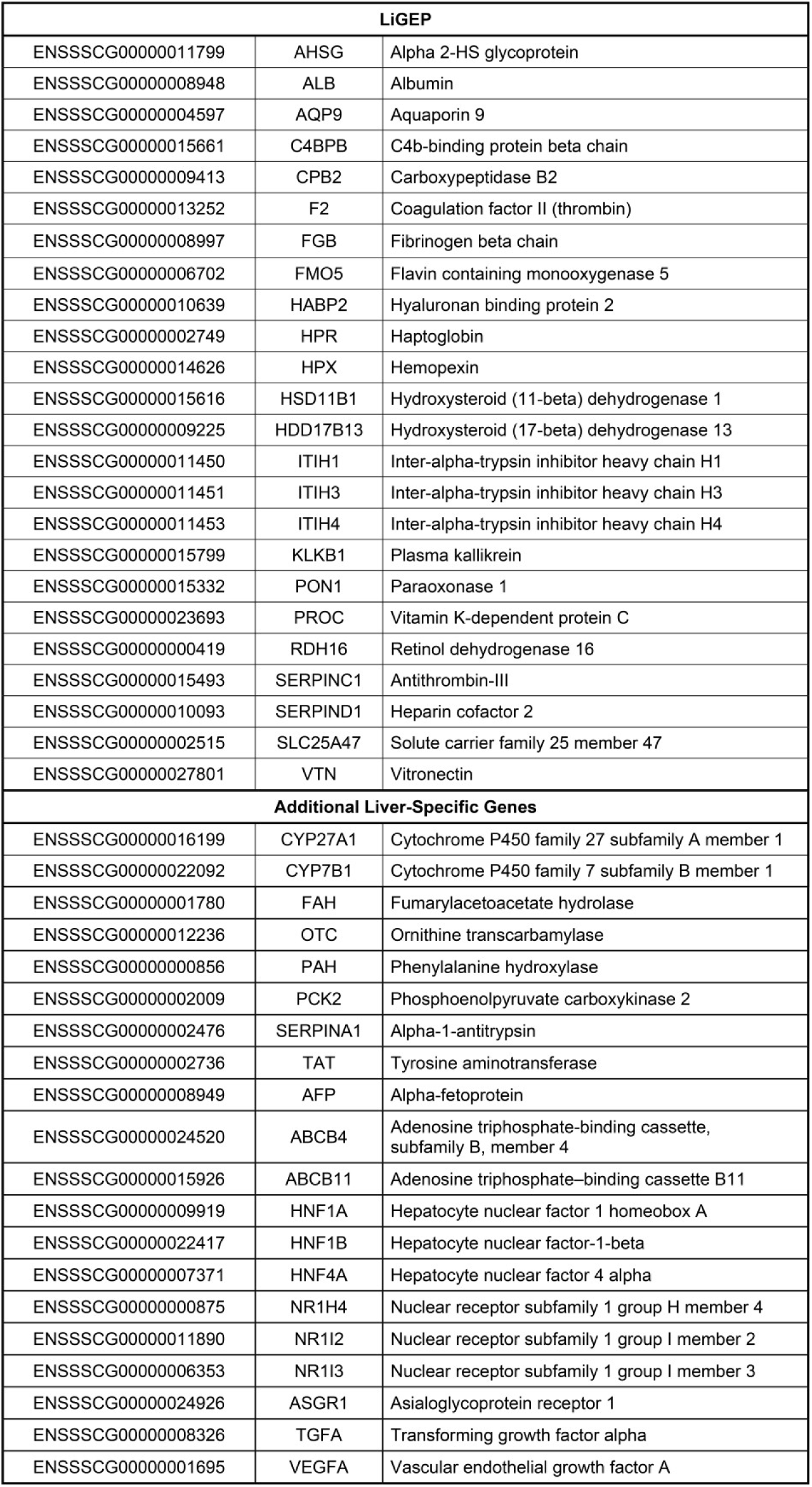
Liver-specific genes included in RNA sequencing analysis

**Supplementary Figure 4.**
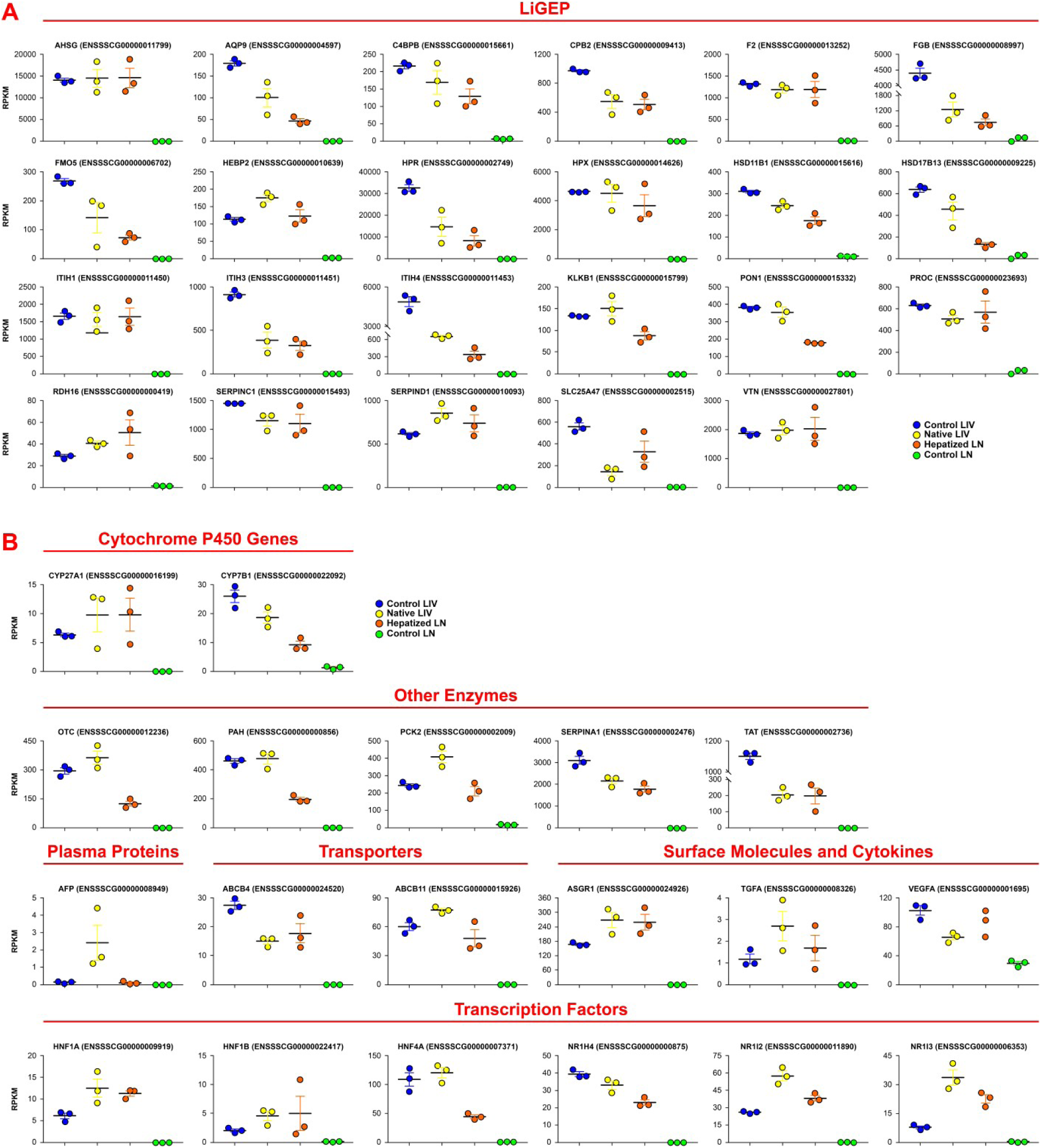
Further RNA sequencing analysis. Scatter dot plot graphs showing RPKM values for LiGEP (liver-specific gene expression panel) (A) or additional liver-specific genes (B) in control liver, native liver, hepatized lymph node, and control lymph node. Data are mean ± SEM.

Supplementary Video 1. 3D render movie of PET-CT images of ^89^Zr-labeled hepatocytes at 6 h post-transplantation into mesenteric lymph nodes in pig.

Supplementary Video 2. 3D render movie of PET-CT images of NIS-labeled hepatocytes at 3 months post-transplantation into mesenteric lymph nodes in pig 265.

Supplementary Video 3. 3D render movie of PET-CT images of NIS-labeled hepatocytes at 6 months post-transplantation into mesenteric lymph nodes in pig 265.

Supplementary Video 4. 3D render movie of PET-CT images of NIS-labeled hepatocytes at 5 months post-transplantation into mesenteric lymph nodes in pig 268.

Supplementary Video 5. 3D render movie of PET-CT images of NIS-labeled hepatocytes at 6 months post-transplantation into mesenteric lymph nodes in pig 268.

